# Region-specific homeostatic identity of astrocytes is essential for defining their reactive phenotypes following pathological insults

**DOI:** 10.1101/2023.02.01.526708

**Authors:** Natallia Makarava, Olga Mychko, Kara Molesworth, Jennifer Chen-Yu Chang, Rebecca J. Henry, Natalya Tsymbalyuk, Volodymyr Gerzanich, J. Marc Simard, David J. Loane, Ilia V. Baskakov

## Abstract

The transformation of astrocytes into reactive states constitutes a biological response of the central nervous system under a variety of pathological insults. Astrocytes display diverse homeostatic identities, which are developmentally predetermined and regionally specified. Upon transformation into reactive states associated with neurodegenerative diseases and other neurological disorders, astrocytes acquire diverse reactive phenotypes. However, it is not clear whether their reactive phenotypes are dictated by regionspecific homeostatic identity or, alternatively, by the nature of an insult. To address this question, regionspecific gene expression profiling was performed for four brain regions (cortex, hippocampus, thalamus and hypothalamus) in mice using a custom Nanostring panel consisting of selected sets of genes that report on astrocyte functions and their reactivity for five conditions: prion disease, traumatic brain injury, brain ischemia, 5XFAD Alzheimer’s disease model and normal aging. Upon transformation into reactive states, genes that are associated predominantly with astrocytes were found to preserve region-specific signatures suggesting that they respond to insults in a region-specific manner. A common gene set was found to be involved in astrocyte remodeling across insults and normal aging. Regardless of the nature of an insult or insult-specificity of astrocyte response, strong correlations between the degree of astrocyte reactivity and perturbations in their homeostasis-associated genes were observed within each individual brain region. The insult-specific populations did not separate well from each other and instead partially overlapped, forming continuums of phenotypes. The current study demonstrates that astrocytes acquire their reactive phenotypes according to their region-specific homeostatic identities. Within these region-specified identities, reactive phenotypes show continuums of states, partially overlapping between individual insults.

## Introduction

The transformation of astrocytes into reactive states constitutes a biological response of the central nervous system (CNS) under a variety of pathological conditions including traumatic brain injury (TBI), ischemic stroke, brain infection and neurodegenerative diseases such as Alzheimer’s, Parkinson’s and prion diseases (Phatnani and Maniatis, 2015; Ferrer, 2017). In contrast to microglia, whose primary function is to protect the brain from pathogens and to clear debris, astrocyte ability to respond to a variety of pathological conditions is one among their many other vital functions. Among these functions are support of neuronal growth, modulation of neurotransmission, formation and maintenance of synapses, regulation of blood flow, maintenance of the blood-brain-barrier (BBB), and supply of energy and metabolic support to neurons, and more (Sofroniew and Vinters, 2010; Dallérac et al., 2018; Santello et al., 2019). It is not clear whether in their reactive state, these critical astrocytic functions become universally affected, and whether transition into reactive state adds to brain impairments or promotes neuroprotection and neuronal repair.

Studies that employed imaging, physiological and transcriptome approaches revealed that under normal conditions, astrocytes are functionally diverse and form distinct neural-circuit-specialized subpopulations (Ben Haim and Rowitch, 2017; Chai et al., 2017; Morel et al., 2017; Zeisel et al., 2018; Khakh and Deneen, 2019; Miller et al., 2019; Batiuk et al., 2020; Huang et al., 2020). In fact, seven developmentally predetermined sub-types of astrocytes that reside in different brain regions have been identified in mouse brains (Zeisel et al., 2018). Upon transformation into reactive states associated with neurodegenerative diseases and other neurological disorders, astrocytes also acquire diverse and sometimes mixed phenotypes (Zamanian et al., 2012; Liddelow and Barres, 2017; Habib et al., 2020; Sofroniew, 2020; Wheeler et al., 2020; Baskakov, 2021; Jiwaji et al., 2022). However, the role of a region-specific homeostatic identity of astrocytes versus a nature of an insult in defining their reactive phenotypes is not well understood. Do astrocytes respond to pathological insults in an insult-specific manner adopting insult-specific states? If so, are insult-specific states uniform across brain regions? Investigating the role of the homeostatic identity of astrocytes versus the nature of pathological insult in driving their reactive phenotypes is important for learning how astrocyte reactivity can be modulated to alleviate pathology.

In order to define the role of regional identity versus nature of insult, the current study performed gene expression profiling under five conditions: prion disease, TBI, brain ischemia, 5XFAD Alzheimer’s disease (AD) model (Oakley et al., 2006), and normal aging (24 month old). We opted for a targeted approach and a bulk tissue analysis using a custom Nanostring panel consisting of selected sets of genes that report on astrocyte functions and their reactivity, and collected data from four brain region: cortex, hippocampus, thalamus and hypothalamus. We found that astrocytes acquire their reactive phenotypes according to their region-specific homeostatic identities. Within region-specified identities, reactive phenotypes represent continuums of activation states, partially overlapping between individual insults.

## Materials and Methods

### Ethics approval and consent to participate

The study was carried out in accordance with the recommendations in the Guide for the Care and Use of Laboratory Animals of the National Institutes of Health. The animal protocol was approved by the Institutional Animal Care and Use Committee of the University of Maryland, Baltimore (Assurance Number: A32000-01; Permit Number: 0118001).

### Prion infection

Using isoflurane anesthesia, 6-week-old C57Bl/6J female and male mice were subjected to intraperitoneal inoculation of 1% brain homogenate, prepared in PBS (pH 7.4) using terminally ill SSLOW-infected mice. Brain materials from the 5^th^ passage of SSLOW in mice was used for inoculations (Makarava et al., 2012; Makarava et al., 2020a). Inoculation volume was 200 μl. 1xPBS was inoculated into normal control groups (Norm). Animals were regularly observed for signs of neurological impairment: abnormal gate, hind limb clasping, lethargy, and weight loss. Mice were considered terminally ill when they were unable to rear and/or lost 20% of their weight.

### Traumatic brain injury

Traumatic brain injury was administered as a Controlled Cortical Impact (CCI) using a custom-designed CCI device, consisting of a microprocessor-controlled pneumatic impactor with a 3.5 mm diameter tip as described (Henry et al., 2020). Briefly, mice were anesthetized with isoflurane administered through a nose mask, and placed on a heated pad to maintain their core body temperature at 37°C. The head was mounted in a stereotaxic frame, a 10-mm midline incision was made over the skull and the skin and fascia were reflected. A 5-mm craniotomy was made on the central aspect of the left temporoparietal bone, between the bregma and lambda. The impounder tip of the injury device was then extended to its full stroke distance (44 mm), positioned to the surface of the exposed dura, and reset to impact the cortical surface. Moderate-level CCI was induced using an impactor velocity of 6 m/s, with a deformation depth of 2 mm as described (Henry et al., 2020). After injury, the incision was closed with interrupted 6-0 silk sutures, anesthesia was terminated, and the animal was placed into a heated chamber to maintain normal core temperature for 45 minutes post-injury. Control mice underwent sham procedure consisting of anesthesia, skin reflection and suture, but did not receive the craniotomy or impact. Mice were monitored daily post-injury.

### Transient middle cerebral artery occlusion (MCAO)

The procedure for MCAO in mice has been described in detail (Bertrand et al., 2017). Briefly, mice (20-25 gm, 8–12 weeks) were anesthetized [induction, 3.0% isoflurane; maintenance, 1.5–2.0% isoflurane with a mixture of O_2_ (200 mL/min) and N2O (800 mL/min)]. Body temperature is maintained during surgery (36.5 ± 0.5 °C) with a feedback heating-controlled pad system (Harvard Apparatus). A midline ventral neck incision was made, and the left common carotid artery (CCA) was ligated proximal to the bifurcation. A second suture was placed around the CCA near the bifurcation. A small arteriotomy was made in the CCA between two sutures. A silicon filament (602356PK5Re Doccol Corp, Redlands CA) was introduced into CCA and advanced into the ICA, ~8.0 mm from bifurcation until it occludes the MCA, where resistance is felt. The silicon filament was secured by sutures, and is left in place for 2 hours. Neurological behavior was monitored, and any animal that does not show circling behavior was excluded. After 2 hours, the animal was re-anesthetized, the occluder filament was removed, and the CCA was ligated.

### Brain tissue collection and RNA isolation

At the designated time points (Table S1), mice were euthanized by CO_2_ asphyxiation and their brains were immediately extracted. The brains were kept ice-cold for prompt dissection or preserved in 10% buffered formalin (MilliporeSigma) for histopathology.

Extracted ice-cold brains were dissected using a rodent brain slicer matrix (Zivic Instruments, Pittsburg, PA). A 2 mm central coronal section of each brain was used to collect individual regions. Allen Brain Atlas digital portal (http://mouse.brain-map.org/static/atlas) was used as a reference. The hypothalami (HTh), as well as thalami (Th), hippocampi (Hp) and cortices (Ctx) were collected into RNase-free, sterile tubes, frozen in liquid nitrogen and stored at −80°C until the RNA isolation with Aurum Total RNA Mini Kit (Bio-Rad, Hercules, CA, USA), as described (Makarava et al., 2020b). Brain tissue samples were homogenized within RNase-free 1.5 ml tubes in 200 μl of Trizol (Thermo Fisher Scientific, Waltham, MA, USA), using RNase-free disposable pestles (Fisher scientific, Hampton, NH). After homogenization, an additional 600 μl of Trizol was added to each homogenate, and the samples were centrifuged at 11,400 x g for 5 min at 4°C. The supernatant was collected, incubated for 5 min at room temperature, then supplemented with 160 μl of cold chloroform and vigorously shaken for 30 sec by hand. After an additional 5 min incubation at room temperature, the samples were centrifuged at 11,400 x g for 15 min at 4°C. The top layer was transferred to new RNase-free tubes and mixed with an equal amount of 70% ethanol. Subsequent steps were performed using an Aurum Total RNA Mini Kit (Bio-Rad, Hercules, CA, USA) following the manufacturer instructions. Isolated total RNA was subjected to DNase I digestion. RNA purity and concentrations were estimated using a NanoDrop One Spectrophotometer (Thermo Fisher Scientific, Waltham, MA, USA).

### Design of Nanostring nCounter Mouse Astrocyte Panel

To design a custom-based nCounter Mouse Astrocyte Panel, publicly accessible databases (www.brainrnaseq.org and https://www.networkglia.eu/en/astrocyte) were used for selecting 275 genes that under normal conditions, express primarily in astrocytes and were assigned to functional pathways. In addition, 47 genes reporting on reactive phenotypes, including A1-, A2-, pan-specific makers and other markers of reactive astrocytes were used for the panel, as well as 8 microglia-, 10 neuron-, and 2 oligodendrocyte-specific genes. The addition of 10 housekeeping genes completed the panel bringing the total number of genes to 352. These genes were assigned to 23 gene sets to allow for an advanced analysis of their group changes (Table S2).

### Analysis of gene expression by Nanostring

Samples were processed by the Institute for Genome Sciences at the University of Maryland, School of Medicine using a custom nCounter Mouse Astrocyte Panel. Only samples with an RNA integrity number RIN >7.2 were used for Nanostring analysis. All data passed QC, with no imaging, binding, positive control, or CodeSet content normalization flags. The analysis of data was performed using nSolver Analysis Software 4.0, including nCounter Advanced Analysis (version 2.0.115) for Principal Component Analysis (PCA) and differential expression analysis. For agglomerative clustering and heat maps, genes with less than 10% of samples above 20 counts were excluded. Z-score transformation was performed for genes. Clustering was done using Euclidian distance and the linkage method was Average.

Differentially expressed genes (DEGs) were calculated using NanoString nCounter Advanced Analysis, for each brain region separately. Each experimental group was compared to the corresponding control group with all other groups present in the analysis. Only genes with adjusted *p*<0.1, and linear fold change ≥±1.2 were counted.

Undirected global significance scores, which measure the cumulative evidence for the differential expression of genes in a pathway, were obtained during Gene Set Analysis performed for all sample groups in parallel (GSA scores) using nCounter built-in algorithms. Each brain region was analyzed separately. To characterize the relationship between astrocyte reactivity and function, GSA scores of corresponding gene sets were summed and plotted against each other as in Figure 2, or further normalized by control group GSAs and plotted as in Figure 6B.

### Histopathology and Immunofluorescence

Formalin-fixed prion-infected brains and mock-inoculated controls were treated for 1 hour with 95% formic acid to deactivate prion infectivity before being embedded in paraffin. All other brains were embedded without formic acid treatment. Subsequently, 4 μm brain sections produced using a Leica RM2235 microtome were mounted on slides and processed for immunohistochemistry. To expose epitopes, slides were subjected to 20 min of hydrated autoclaving at 121°C in Antigen Retriever citrate buffer, pH 6.0 (C9999, MilliporeSigma). For co-immunofluorescence, rabbit polyclonal anti-Iba1 antibody was used in combination with chicken polyclonal anti-GFAP antibody. The secondary antibodies were goat anti–rabbit or anti-chicken IgG conjugated with Alexa Fluor 546 for red color or Alexa Fluor 488 for green color (Thermo Fisher Scientific). Images were collected with an inverted microscope (Nikon Eclipse TE2000-U) equipped with an illumination system X-cite 120 (EXFO Photonics Solutions Inc., Exton, PA, United States) and a cooled 12-bit CoolSnap HQ CCD camera (Photometrics, Tucson, AZ, United States). Fiji ImageJ software was used for image processing.

### Statistical analysis

Differential expression analysis was performed with NanoString nCounter Advanced Analysis software. Only genes with adjusted *p*<0.1, and linear fold change ?±1.2 were counted as Differentially Expressed Genes (DEGs). Adjustment of *p*-values was done with the Benjamini-Yekutieli method. For Figure S6, normalized counts for individual genes were calculated and plotted as Mean ± Standard Deviation using GraphPad Prism 9.2.0. *P*-values for the differences between experimental samples and their corresponding controls, if significant, are shown in the Supplementary Table S3. The number of individual samples in each group is provided in Supplementary.

### Key resources

| REAGENTS | SOURCE | IDENTIFIER |
| --- | --- | --- |
| **Antibodies** |  |  |
| Anti-Iba1, rabbit polyclonal | FUJIFILM Wako | #013-27691 |
|  | Pure Chemical Corporation |  |
| Anti-GFAP, chicken polyclonal | Millipore Sigma | #AB5541 |
| Goat anti-Rabbit, Alexa Fluor 546 | Thermo Fisher Scientific | #A-11035 |
| Goat anti-Chicken, Alexa Fluor 488 | Thermo Fisher Scientific | #A-11039 |
| **Chemicals** |  |  |
| DAPI | Thermo Fisher Scientific | #62248 |
| PBS, pH7.4 | Thermo Fisher Scientific | #AM9624 |
| 10% buffered formalin | Millipore Sigma | #HT501128 |
| Trizol | Thermo Fisher Scientific | #15596026 |
| Chloroform | Millipore Sigma | #C2432 |
| Ethanol (for RNA extraction) | Millipore Sigma | #E7023 |
| Formic acid | Millipore Sigma | #F0507 |
| Citrate Buffer, pH 6.0, Antigen Retriever | Millipore Sigma | #C9999 |
| **Critical commercial assays** |  |  |
| Aurum Total RNA Mini Kit | Bio-Rad | #7326820 |
| Custom nCounter Mouse Astrocyte Panel | NanoString Technologies | UM_Astrocyte_1_C6977 |
| **Experimental models** |  |  |
| Mouse adapted prion strain SSLOW | Makarava et al., J Clin Invest., 2020 | N/A |
| 5XFAD (B6 SJLF1/J) | The Jackson Lab | JAX MMRRC Stock# 034840 |
| **Software and algorithms** |  |  |
| nSolver Analysis Software 4.0 | NanoString Technologies | https://www.nanostring.com/products/analysis-solutions/ncounter-analysis-solutions/ |
| nCounter Advanced Analysis (v. 2.0.115) | NanoString Technologies | https://www.nanostring.com/products/analysis-solutions/ncounter-advanced-analysis-software/ |
| GraphPad Prism 9.2.0 | GraphPad Software Inc | https://www.graphpad.com/scientific-software/prism/ |
| Fiji ImageJ 1.53c | NIH | https://imagej.net/Fiji |

### Data availability

All data generated or analyzed during this study are included in this published article and its supplementary information file. Mouse-adapted prion strain SSLOW used in this study is available from the lead contact, but may require a completed materials transfer agreement.

## Results

### Astrocyte panel detects changes in gene expression profiles upon various pathological insults

For analysis of gene expression, we designed a custom Nanostring panel that reports on astrocyte function genes and reactive state (Makarava et al., 2021). The panel consisted of 275 genes, which were previously found to capture astrocyte homeostatic functions. An additional 47 genes were selected to report on astrocyte reactivity including A1-, A2- and pan-specific makers along with other markers of reactive astrocytes and genes that regulate astrocyte reactivity. In addition, the panel also included 8 microglia-, 10 neuron-, and 2 oligodendrocyte-specific genes along with 10 housekeeping genes. Gene expression was analyzed for five experimental conditions: prion infection, TBI, ischemic stroke, Aβ plaque formation, and normal aging (Table S1). For each condition, datasets were collected for four brain regions - cortex (Ctx), hippocampus (Hp), thalamus (Th), and hypothalamus (HTh) (Table S2).

For infecting animals with prions, we used mouse-adapted strain SSLOW, which has the shortest incubation time to disease among mouse-adapted prion strains and induces profound widespread neuroinflammation across brain regions (Fig. S1A, C and E; Table S3) (Makarava et al., 2020a; Makarava et al., 2021). Male and female C57Bl/6J mice were infected with SSLOW via intraperitoneal route (Table S1).

As a model of TBI, we employed the Closed Cortical Impact model in adult female C57Bl/6J mice, and harvested ipsilateral and contralateral brain tissues 7 and 60 days after the primary insult (Table S1), except for hypothalamus, which was harvested as one sample containing both ipsilateral and contralateral parts. Consistent with the sustained injury, the gene expression profile of the ipsilateral cortex was significantly altered in comparison to the cortex of sham control animals (Fig. S2A). Interestingly, the ipsilateral hippocampus and thalamus displayed even more profound changes in gene expression than the cortex (Fig. S2A; Table S3). In contralateral regions of the brain, only minimal gene expression changes were detectable. Many genes remained upregulated 60 days post-injury, but to a lesser degree in comparison to the 7 DPI (days post-injury) group (Fig. S2A; Table S3). Thus, for comparison with other insults, we chose the ipsilateral samples obtained 7 days after injury, which is the peak of post-traumatic neuroinflammation in this model (Loane et al., 2014; Villapol et al., 2017).

As a model of ischemic stroke, we used Middle Cerebral Artery Occlusion (MCAO) in adult female C57Bl/6J mice (Table S1). The blood flow was occluded for 1 hour, followed by 24 hours or 5 days of reperfusion, at which point the animals were euthanized. Since brain tissue was collected from a narrow 2 mm coronal section, and due to a possible variation in the location or the extent of the stroke, MCAO samples displayed a higher degree of variability in comparison with the other insults (Fig. S2B; Table S3). We chose to focus on the 24-hour time point post-ischemia, which displayed a milder response but more consistent changes relative to the 5-day samples (Fig. S2B).

To assess the changes in the gene expression related to AD, we employed the 5XFAD mouse model and harvested brain tissue from 10-month-old male and female mice. These mice carry five mutations associated with a familial form of AD, and at 10 months of age, they display behavioral deficits and a full spectrum of pathological changes including mature amyloid plaques, profound neuroinflammation and significant changes in gene expression (Fig. S1G, H and S2C; Table S3) (Oakley et al., 2006). Age-matched littermates that lacked disease-associated variants (B6SJLF1/J, referred to as WT) were used for comparison (Table S1). As detected by heat map analysis (Fig. S2C), female 5XFAD mice had somewhat stronger changes in gene expression, which was in good agreement with published reports of higher plaque burden in 5XFAD females in comparison to males (Oakley et al., 2006).

To analyze gene expression changes upon aging, 24-month-old C57Bl/6J male and female mice were compared with C57Bl/6J control animals of younger age: 3 and 5-month-old controls for TBI (n=6), and normal 8 to 13-month-old mice (n=9) (Table S1). A heat map of normalized counts for individual samples visualized the level of variations between age cohorts (Fig. S2D). Ultimately, the youngest animals in the study, 3-month-old mice that correspond to mature adults, were used as a reference in differential expression analysis of aged mice (24-month-old).

### Region-specificity of astrocyte gene expression is maintained after insults

Individual samples within control groups demonstrated robust reproducibility and high regional specificity in their gene expression pattern (Fig. 1A and S3). Indeed, regardless of animal age, sex or strain of mice, all control samples clustered in a strict accordance with brain region showing only minor variations between different control groups (Fig. S3). These data were consistent with the previous reports that documented well-defined regionspecific homeostatic identities of astrocytes (Batiuk et al., 2020; Bayraktar et al., 2020; Habib et al., 2020; Westergard and Rothstein, 2020; Makarava et al., 2021). Remarkably, in all pathological insults, the transformation of astrocytes into disease-associated states occurred in a region-specific manner, as astrocytes preserved region-specific identities in their disease-associated states (Fig. 1A and S4). Preservation of regional identities by reactive astrocytes contrasts with the transformation of microglia, which acquire a more uniform disease-associated signature across brain regions (Makarava et al., 2020b).

**Figure 1.**
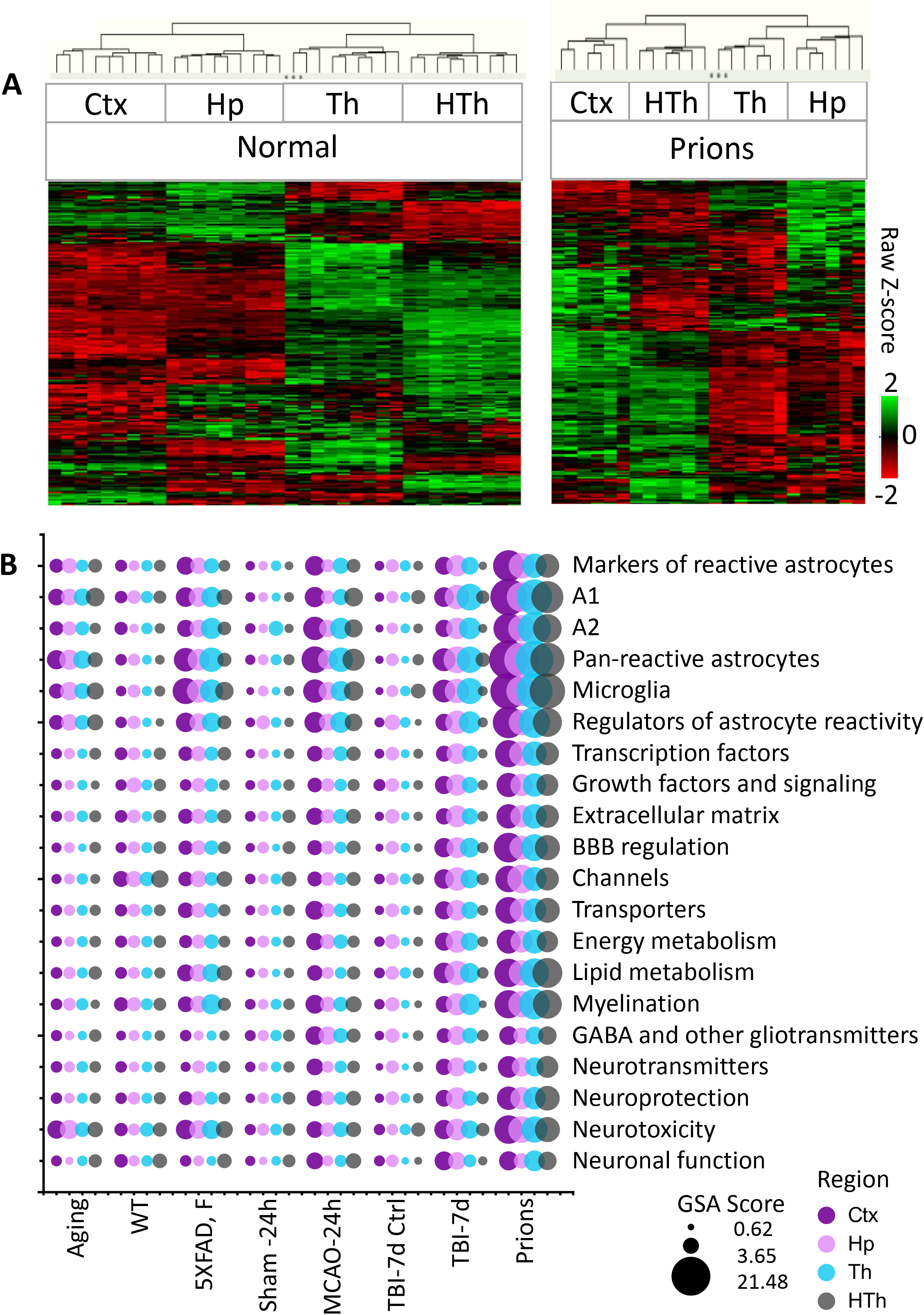
Analysis of gene expression in different brain regions. **(A)** Hierarchical clustering of mock-inoculated samples from Normal group (left) and Prions group (right) showing high reproducibility of the Astrocyte panel and regional specificity of astrocyte function genes, which is maintained at the terminal stage of prion infection. Clustering was performed using genes related to BBB regulation, lipid and energy metabolism, extracellular matrix, junction, myelination, channels, transporters, gliotransmitters and neurotransmitters, neuroprotection and neurotoxicity, and astrocyte-specific markers (Table S2). **(B)** Bubble Plot of undirected global significance scores obtained with Nanostrong Gene Set Analysis (GSA) by comparison of all groups with Normal samples (Table S1). TBI-7d is represented by ipsilateral Ctx, Hp and Th, and whole HTh. The size of the bubbles reflects the degree of regional changes for individual gene sets with different insults. Ctx – cortex, Hp – hippocampus, Th – thalamus, HTh – hypothalamus.

**Figure 2.**
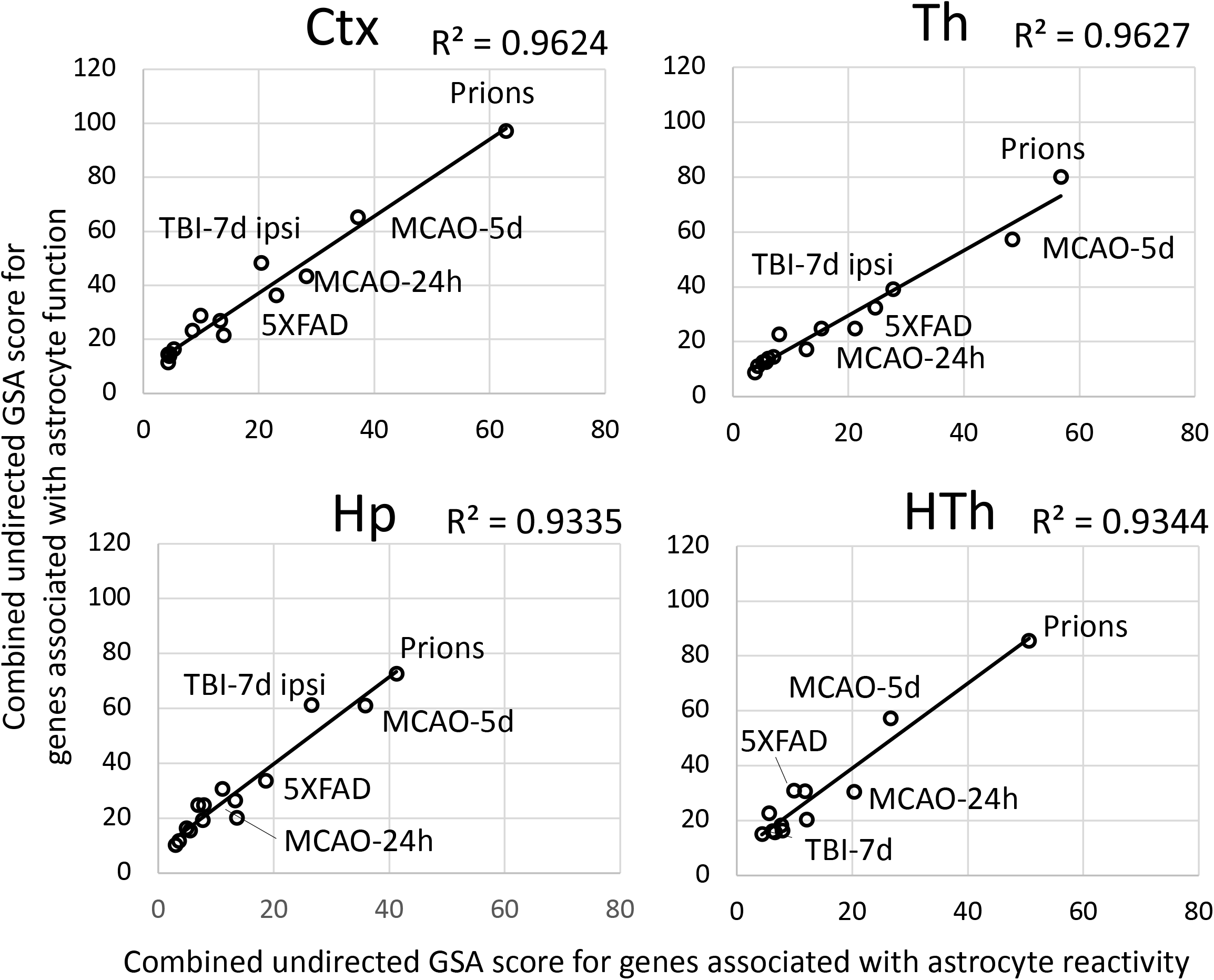
Relationship between the changes in the expression of gene sets in different brain regions. A strong correlation between the sum of undirected global significance scores of the gene sets reporting on the degree of astrocyte reactivity (A1-, A2-, pan-specific markers and other markers of reactive astrocytes) and dysregulation of astrocyte function (BBB regulation, channels, lipid metabolism, myelination, energy metabolism, extracellular matrix, gliotransmitters, neurotransmitters, transporters, neuroprotection, neurotoxicity). Ctx – cortex, Hp – hippocampus, Th – thalamus, HTh – hypothalamus.

### The degree of astrocyte reactivity correlates with the degree of changes in genes associated with astrocyte functions

Within individual regions, the degree of astrocyte reactivity varied depending on the insult (Fig. 1B and S5). Regardless of the brain region, the highest gene set analysis (GSA) scores that reported on astrocyte reactivity and functional changes were found in prion-infected animals (Fig. 1B and S5). Plotting combined undirected GSA scores that reflect astrocyte reactivity against the combined undirected GSA scores for physiological functions revealed a very strong correlation between the two (Fig. 2). Remarkably, as judged from the R^2^ values, the strength of the relationship was very high in all four regions examined (Fig. 2). These results suggest that regardless of the nature of the insult or the identity of astrocyte reactive phenotype, the changes in functional pathways are tightly linked to and, perhaps, determined by the degree of astrocyte reactivity.

### Region-specific continuums of astrocyte reactive states

To analyze region-specific insult-elicited differences across all groups, we performed a Principal Component Analysis (PCA) which placed the individual samples of different insults into continuums of phenotypes (Fig. 3). PCA of an entire panel resulted in two well-resolved continuums: one shared by cortex and hippocampus, and another shared by thalamus and hypothalamus (Fig. 3A, left panels). Within each continuum, overlaps between individual insult-dictated phenotypes were observed (Fig. 3A, right panels). As expected, phenotypes in normal controls overlapped substantially with those of Aging group. Furthermore, Aging overlapped with 5XFAD and MCAO-24h, 5XFAD and MCAO-24h overlapped with TBI-7d, and finally, TBI-7d with prions. Similar patterns were observed for smaller gene sets that report on specific functions, including Neuroprotection/Neurotoxicity, Transporters, Metabolism, Reactive Astrocytes, and others (Fig. 3B). However, depending on the gene set employed, a better separation of cortex-versus hippocampus-specific continuums (Metabolism) or thalamus-versus hypothalamus-specific continuums (Transporters, Neuroprotection/Neurotoxicity) can be seen (Fig. 3B).

**Figure 3.**
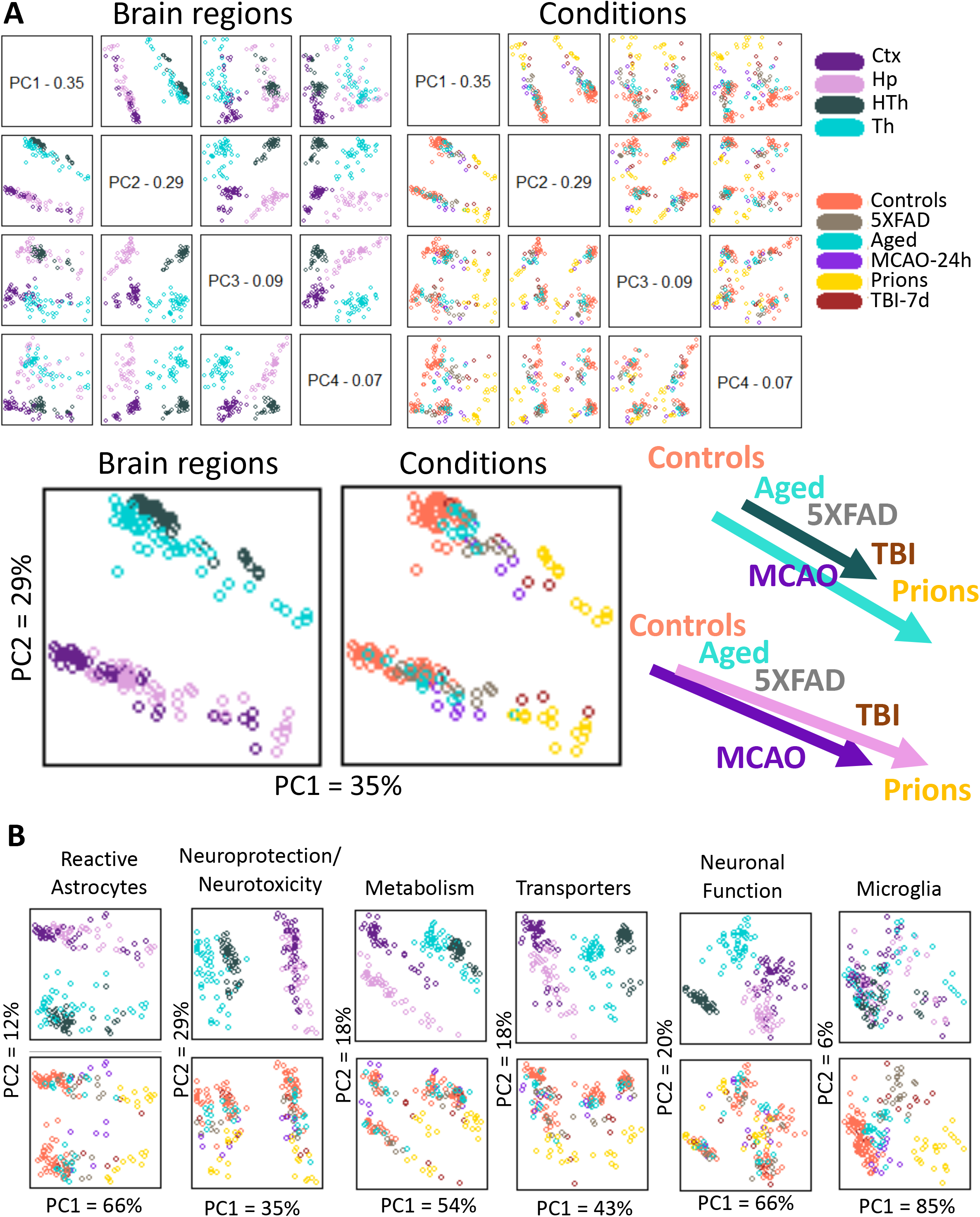
Principle Component Analysis. **(A)** Principle Component Analysis (PCA) performed with the entire panel of genes for samples from 5 experimental groups and 4 brain regions. Control group in PCA included Normal, TBI-7d Ctrl, and Sham-24h samples combined. Distribution of samples according to brain region and to condition is shown on the top left and top right, respectively. Enlarged PC1 vs PC2 plots are shown below, with a schematic illustrating continuums of astrocytic phenotypes on the right. **(B)** PCA performed with selected gene sets for the same samples. Each point in PCA represents individual animal. Ctx – cortex, Hp – hippocampus, Th – thalamus, HTh – hypothalamus.

To summarize, (i) astrocyte phenotypes associated with different pathological insults partially overlapped and, together, constituted continuums of states; (ii) individual continuums were region-specific. In comparison to astrocytes, microglia did not form well-defined region-specified phenotypes, showing mostly insult-specific differences, which were partially overlapping with an exception of prions (Fig. 3B). Gene sets reporting on neuronal functions displayed an opposite pattern: well-defined separation according to brain region, but weak insult-specific separation within individual regions (Fig. 3B).

### Astrocyte remodeling involves a core gene set common across insults

A comparison of differentially expressed genes (DEGs) detected a significant overlap in DEGs between experimental conditions including normal aging (Fig. 4A, Table S3), suggesting that the same core gene set is involved in remodeling regardless of insult. Of particular interest is the Aged group that shared a significant number of DEGs with each studied insult (Fig. 4B; Table S4). Out of 32 DEGs identified in the cortex of the Aged cohort, the number of DEGs shared with other conditions increased from 53% for 5XFAD and 56 % for MCAO-24h to 81% for TBI-7d and prion groups. On top of the upregulated DEGs of the core gene set, which is defined as a set dysregulated across all conditions including Aging, were markers of reactive astrocytes GFAP, Vim and Serpina3n, microglia markers CD68 and TLR2, and complement factor C4a. In addition, gene related to astrocyte function Slc14a1 (urea transporter) was within the core gene set (Table S4).

**Figure 4.**
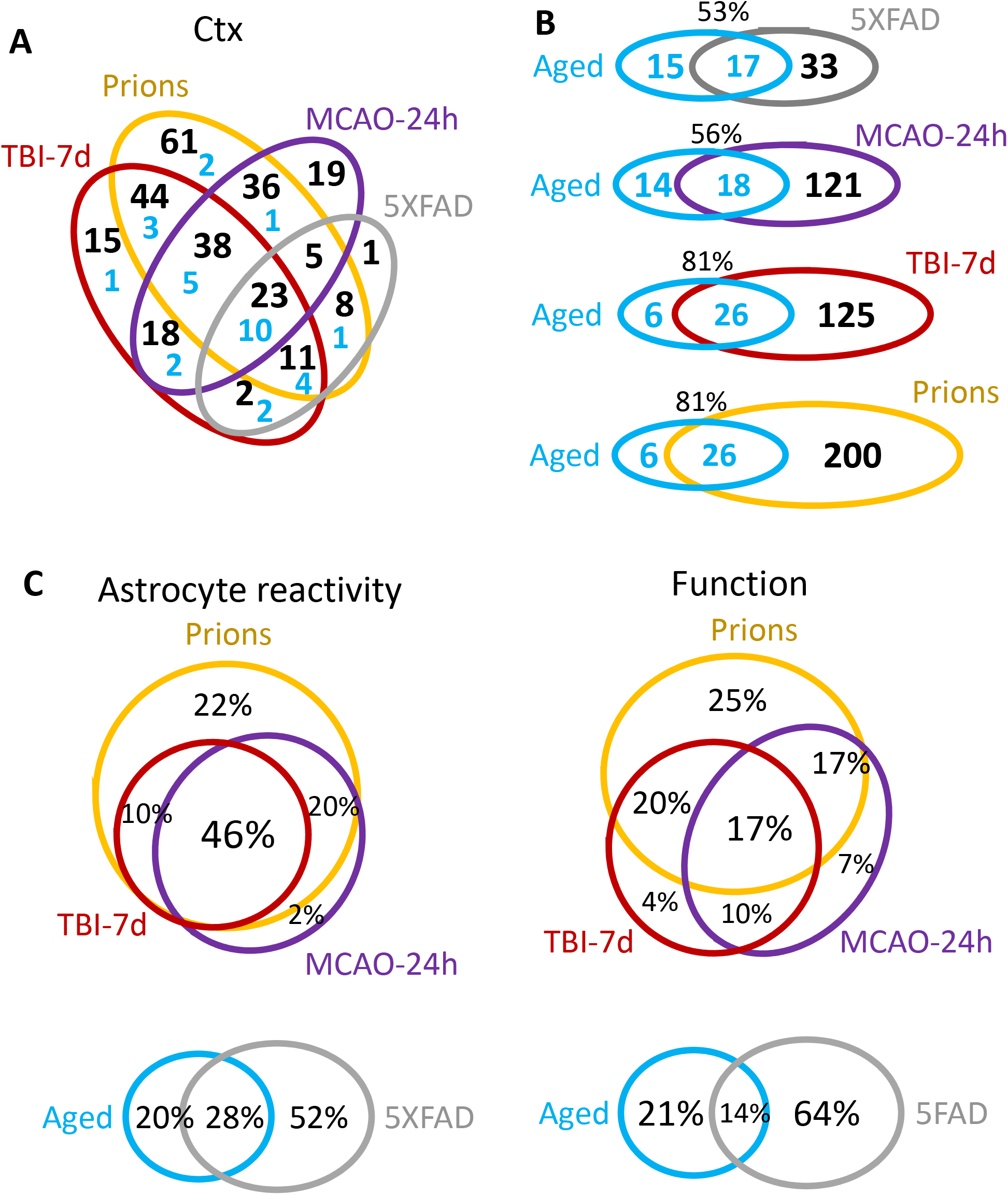
Venn diagrams of differentially expressed genes in cortex. **(A)** Venn diagram for DEGs in the cortex of 5XFAD (females and males, n=3+3), MCAO-24h (females, n=3), TBI-7d (females, n=3), and Prions (females and males, n=3+3). Blue numbers represent a portion of DEGs common with the Aged group (females and males, n=3+3). **(B)** Venn diagrams representing overlaps in DEGs detected for Aged cortex and for cortexes of other groups. **(C)** Venn diagrams of DEG percentage overlap in the reporters of astrocyte reactivity (left) and function (right), presented for the groups with strong changes in gene expression (top) and mild changes in gene expression (bottom). Genes annotated as A1, A2, pan-reactive, other markers of reactive astrocytes, regulators of astrocyte reactivity and astrocyte-specific markers were considered as reporters of Inflammation. Genes related to BBB regulation, lipid and energy metabolism, extracellular matrix, junction, myelination, channels, transporters, gliotransmitters and neurotransmitters, and neuroprotection were included into Function DEGs. Differential expression analysis was performed using Nanostring Advanced Analysis software. Each group was compared to their corresponding controls (Table S1). DEGs for the Aged group were calculated relative to the youngest control group (TBI-7d Ctrl, n=3F). The genes with linear fold greater than 1.2 up or down and adjusted *p*<0.1 were selected as DEGs (Table S3). Ctx – cortex. TBI-7d is represented by ipsilateral Ctx.

### Insult-specific changes in astrocyte phenotypes

The strong correlation between the degree of astrocyte reactivity and the extent of the changes in function genes (Fig. 2) concealed important insult-specific nuances that became evident upon differential expression analysis. The number of detected DEGs in the cortex gradually increased from 32 in Aged and 50 in 5XFAD groups, to 139 in MCAO-24h, 151 in TBI-7d, and 226 in Prions (Table S3). Venn diagrams of cortical DEGs built separately for the markers of reactive state and function-related genes revealed a greater insult-specificity of functional DEGs in comparison to the specificity in reactive marker gene set (Fig. 4C). Remarkably, this trend was observed between groups with strong changes in gene expression (Prions, TBI-7d, MCAO-24h) as wells as groups with mild changes (5XFAD and Aged mice) (Fig. 4C). While the degree of astrocyte reactivity correlates with the extent of dysregulation in genes associated with astrocyte function, the functional changes were not identical between the insults.

Linear fold changes in genes related to homeostatic functions were much lower than the changes in the markers of reactive states (Table S3). Clustering of log2 linear fold changes for all experimental groups revealed that both sets of genes, i.e. those reporting on astrocyte reactivity and functional dysregulation, displayed insult-specific response patterns (Table S3, Fig. 5A and B). For instance, markers of reactive astrocytes GFAP and Serpina3n clustered together, while Vim belonged to a different cluster, pointing to the existence of disease-specific response patterns of astrocyte reactivity (Fig. 5A). The same was true for the set of genes associated with astrocyte functions (Fig. 5B). For example, BBB regulator Agt was detected among DEGs common for Prions and TBI-7d, but it changed in opposite directions: upregulated in TBI-7d, but downregulated in Prions (Fig. S6). MCAO-24h was characterized by a transient downregulation of many genes associated with astrocyte functions (Fig. 5B) that were upregulated in TBI-7d. The expression of the most prevalent water channel Aqp4 increased in cortices of 5XFAD, TBI-7d and Prions, but remained unchanged in MCAO-24h (Fig. S6). Channels and transporters were affected in Prions, TBI-7d and MCAO-24h, but out of these three animal groups, downregulation of Kcna2 and Kcnk1 was accompanied by a statistically significant upregulation of ATPase subunits in the Prions group only (Table S3).

**Figure 5.**
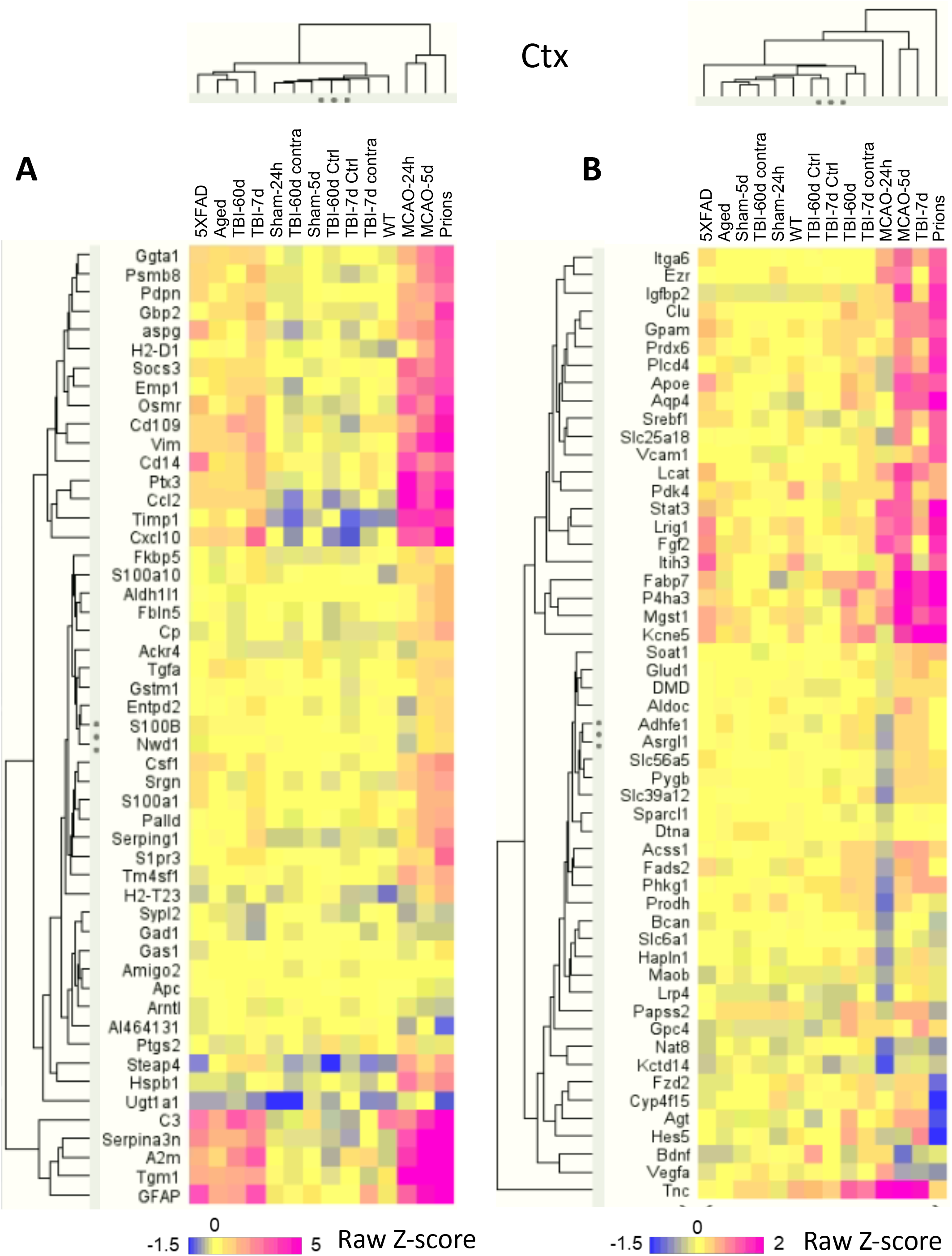
Hierarchical clustering of log2 changes in the expression of selected genes in cortex. **(A)** Selected astrocyte reactivity reporters form several separate clusters with an insult-specific pattern of activation. **(B)** Hierarchical clustering of selected astrocyte function genes highlights insult-specific changes in their expression. Hierarchical clustering was performed for groups with a number of samples as described in Table S1. TBI-7d and TBI-60d are ipsilateral samples. Z-score transformation was performed for genes.

**Figure 6.**
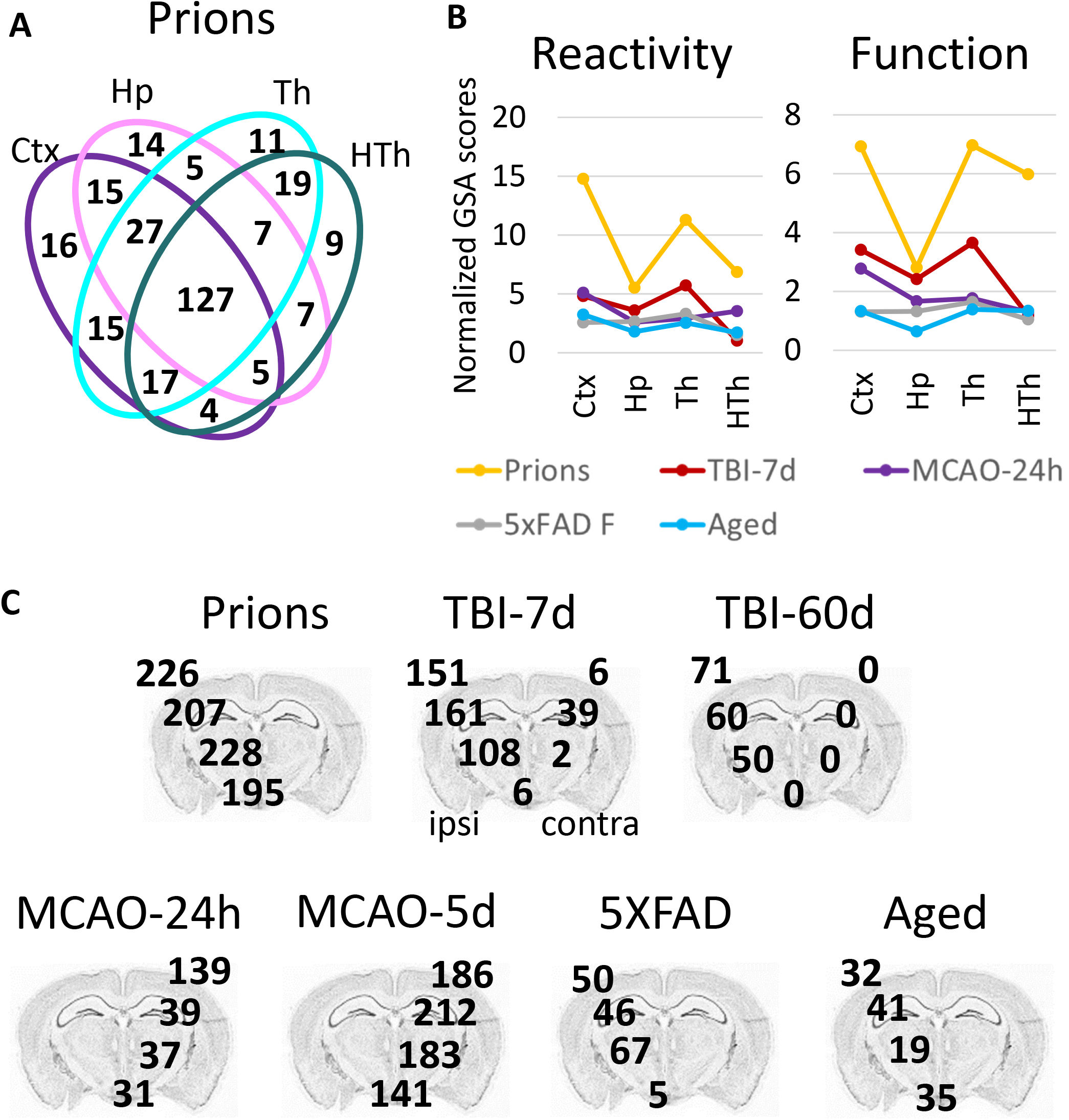
Region-specific changes in gene expression. **(A)** Venn diagram for DEGs in 4 brain regions of Prions group (females and males, n=3+3) calculated in comparison to the Normal group (females and males, n=6+3). **(B)** Combined undirected global significance scores characterizing astrocyte reactivity (left) and function (right) in 4 brain regions upon 5 insults. Undirected global significance scores were calculated with Gene Set Analysis (GSA) for individual gene sets and summed to obtain combined GSA scores for Reactivity and Function for each brain region and insult. Combined GSA scores for Reactivity included GSA scores for A1-, A2-, pan-specific markers and other markers of reactive astrocytes. Combined GSA scores for Function included GSA scores for BBB regulation, lipid and energy metabolism, extracellular matrix, myelination, channels, transporters, gliotransmitters and neurotransmitters, and neuroprotection. To normalize, combined GSA scores for each insult were divided by the corresponding GSA scores obtained for the Normal group. TBI-7d is represented by ipsilateral Ctx, Hp and Th, and whole HTh. **(C)** The number of differentially expressed genes (DEGs) in all experimental groups (Table S1). The numbers for DEGs in cortex, hippocampus, thalamus and hypothalamus are displayed over the corresponding regions of brain. Differential expression analysis was performed using Nanostring Advanced Analysis software. Each group was compared to their corresponding controls (Table S1). DEGs for the Aged group were calculated relative to the youngest control group (TBI-7d Ctrl, n=3F).

To summarize, the response by genes reporting on homeostatic functions displayed a higher degree of disease-specificity relative to the response by markers of reactive astrocytes. Nevertheless, despite insult-specificity in astrocyte response, the degree of net changes in functional gene sets correlated well with the degree of the net changes in astrocyte reactivity (Fig. 2).

### Region-specific changes in astrocytic phenotypes

To study the extent of astrocyte transformations in different brain regions, we focused on cortex, hippocampus, thalamus and hypothalamus in the Prion group. This group induced the strongest gene expression changes throughout the brain, while other insults were preferentially affecting certain regions and/or had a milder impact (Fig. 1B and S5). Despite the fact that region-specificity of astrocytes was maintained even in the Prion group (Fig. 1A), a remarkable overlap of DEGs between Ctx, Hp, Th, and HTh was observed (Fig. 6A). These findings suggested that similar to the insult-specific changes in astrocytic phenotypes, regional transformation involves quantitative changes of a core gene set regardless of brain region.

### Thalamus is vulnerable regardless of the nature of pathological insult

In animals challenged with prion strain SSLOW, prion replication is widespread across the brain (Makarava et al., 2020a; Makarava et al., 2021), whereas other experimental insults used here were expected to target brain regions more selectively. For instance, in TBI and MCAO, the primary impact of injuries were expected to affect the cortex due to the primary injury location and occlusion region. In 5XFAD mice, the earliest and heaviest plaque deposition was found in the cortex and hippocampus (Oakley et al., 2006). Surprisingly, in the current work, the thalamus showed profound astrocyte response in all experimental insults (Fig. 6B). The high vulnerability of thalamus was especially noticeable in TBI, despite the fact that the cortex was the site of injury (Fig. 6B). In TBI, the number of DEGs in the ipsilateral thalamus exceeded those in the contralateral cortex 7 days post insult and remained high even at 60 days post TBI (Fig. 6C). For a comparison, HTh showed only a minimal number of DEGs in TBI-7d (Fig. 6C). In the Prion group and in 5XFAD mice, the number of DEGs in thalamus was higher than in other regions (Fig. 6C). In 5XFAD mice, the number of DEGs in Th exceeded the number of DEGs in Ctx and Hp, while HTh was minimally affected (Fig. 6C). In comparison to animal groups subjected to insults, the natural Aged group showed a mild response of Th with respect to the number of DEGs (Fig. 6C), yet a relatively strong response with respect to GSA score (Fig. 6B). To summarize, the thalamus appears to be a highly vulnerable brain region, reacting not only to a direct insult, but also to injuries delivered to other brain regions.

## Discussion

Studies on single-cell transcriptional profiling revealed that under normal conditions, seven distinct, developmentally predetermined sub-types of astrocytes with well-defined regional specialization exist in the mouse brain (Zeisel et al., 2018; Batiuk et al., 2020). Physiological and morphological studies demonstrated that astrocytes are, indeed, functionally diverse and form distinct neural-circuit-specialized subpopulations in different brain regions (Ben Haim and Rowitch, 2017; Chai et al., 2017; Morel et al., 2017; Zeisel et al., 2018; Khakh and Deneen, 2019; Miller et al., 2019; Batiuk et al., 2020; Huang et al., 2020). Considerably less is known about the phenotypic diversity of astrocytes under pathological conditions (Zamanian et al., 2012; Liddelow and Barres, 2017; Habib et al., 2020; Wheeler et al., 2020). In recent years, several important questions have been under critical discussion. Do astrocytes respond to pathological insults in a specific manner adopting distinct, insult-specific states? Are insult-specific states uniform across brain regions? What are the roles of insult-specific response versus region-specific homeostatic identity in dictating astrocytes reactive phenotype (Escartin et al., 2021)?

To answer these questions, the current studies examined the region-specific response of genes associated with astrocyte functions and reactivity to insults of a diverse nature including prion infection, mechanical injury (TBI), genetic mutations associated with familial AD, ischemic insult and normal aging. We found that upon transformation into reactive states, genes that are associated predominantly with astrocytes preserved a region-specific signatures suggesting that they respond to insults in a region-specific manner. Comparison of the gene sets that report on homeostatic functions with those that report on astrocyte reactivity revealed that they first displayed a higher level of disease-specificity relative to the last. In fact, a common gene set that was found to be involved in astrocyte remodeling across insults and normal aging included markers of astrocyte reactivity (GFAP, Vim, Serpina3n,). Remarkably, regardless of the nature of an insult or insult-specificity of astrocyte response, strong correlations between the degree of astrocyte reactivity and perturbations in their homeostasis-associated genes were observed within each individual brain region. These results suggest that astrocyte reactivity changes in parallel with changes in expression of genes associated with astrocyte functions regardless of the nature of an insult. While PCA did not dismiss the idea regarding the existence of insult-specific phenotypes, the insult-specific populations did not separate well from each other and instead partially overlapped, forming continuums of phenotypes. The continuums of phenotypes were region-specific suggesting that in defining reactive phenotypes, the role of region-specific homeostatic identity perhaps is as important as the nature of an insult. Consistent with the last statement, thalamic astrocytes were found to be the most responsive to insults, as they reacted not only to direct insults within thalamus, but also to injuries delivered to other brain regions.

How many reactive phenotypes of astrocytes exist? Abandoning of the A1 (neurotoxic) /A2 (neuroprotective) binary model renewed the discussion about the diversity of reactive phenotypes (Diaz-Castro et al., 2019; Hartmann et al., 2019; Escartin et al., 2021; Makarava et al., 2021). Previous studies of individual neurodegenerative diseases, including prion disease and multiple sclerosis as well as peripherally induced neuroinflammation, documented that astrocytes respond to insults in a region-specific manner and maintain their region-specific identities (Itoh et al., 2018; Makarava et al., 2019; Wheeler et al., 2020; Diaz-Castro et al., 2021; Makarava et al., 2021). A number of studies focused on the diversity of astrocytes within individual diseases (Diaz-Castro et al., 2019; Al-Dalahmah et al., 2020; Habib et al., 2020; Wheeler et al., 2020), whereas a region-specific profiling of reactive phenotype across multiple diseases has never been performed. Is it the nature of an insult or region-specific homeostatic identity that drives the reactive phenotype? On one hand, within the same brain region, changes in the expression of homeostatic genes showed limited overlaps between individual insults, arguing that the nature of an insult is important in dictating the range of homeostatic functions affected (Fig 4). On the other hand, PCA, which takes into account not only the number of changed genes but also the extent of their changes, revealed a better separation into distinct clusters according to brain region rather than the nature of an insult. For instance, even within the same insult, the phenotypic signatures of cortical and thalami clusters were found to be clearly distinctive (Fig. 3A). Depending on a specific homeostatic gene set, better separations between region-specific signatures of cortex and hippocampus or thalamus and hypothalamus could be achieved (Fig. 3B). Moreover, within the same region, phenotypic signatures for individual insults showed considerable overlaps. Together, these results argue that astrocytes acquire their reactive phenotypes according to the region-specific homeostatic identities. Furthermore, within region-specified identities, reactive phenotypes form continuums of states.

Analysis of the affected physiological pathways suggests that in the reactive phenotypes, global transformation of the homeostatic functions took place, as evident from disturbance in multiple pathways including BBB regulation, transporters, neuroprotection and neurotoxicity, extracellular matrix, myelination, lipid metabolism and others (Fig. S5). Remarkably, the degree of dysregulation in homeostatic functions correlated strongly with the degree of astrocyte reactivity. Are these data purely correlative or a causal relationship between astrocyte reactivity and homeostatic function exists? Within individual brain regions, the relationship between astrocyte reactivity and change in homeostatic functions was strong across insults of different nature, and was not affected by the degree of impact on individual regions. Moreover, regardless of a specific reactive phenotype or function affected by individual insults, the net changes in homeostatic gene sets mirrored the degree of astrocyte reactivity suggesting that the two parameters are tightly coupled and reflective of a global transformation in astrocyte physiology. Whether the net impact of such transformation is beneficial or detrimental is difficult to project from transcriptome analysis alone, and requires functional tests. In previous studies, reactive astrocytes isolated from prion infected animals or mouse models of AD displayed a reduced expression of neuronal support genes and was found to be synaptotoxic in neuronal-astrocyte co-cultures (Orre et al., 2014; Kushwaha et al., 2021).

While the Aged group were not provoked by an insult, in agreement with previous studies (Boisvert et al., 2018; Clarke et al., 2018), their astrocytes displayed clear signs of transformation into reactive phenotypes. Two interesting observations could be made. First, the core gene set that was dysregulated across all of the groups predominantly consisted of genes that report on astrocyte reactivity with only one gene associated with homeostatic functions. This pattern suggests that astrocyte transformation is perhaps driven by changes in gene sets associated with reactivity. Second, the percentage of overlap between DEGs in the Aged group and DEGs in other groups grew with the severity of astrocytic response to individual insults, from 53% overlap with the 5XFAD group to 81% overlap with the Prion group (Fig. 4B). Even though the clusters corresponding to the Prion and the Aged groups were separated far apart in the PCA, which is primarily due to differences in severity of the responses (Fig. 3), the DEGs of the Aged group overlapped almost entirely with the DEGs of the Prion group. This suggests that astrocytic response to prions and other insults engage and perhaps overuse the pathways that are evolved evolutionarily for optimizing astrocyte homeostatic function in normal aging.

Are region-specific populations of astrocytes equally vulnerable to insults? As judged by changes in transcriptome, the rates of astrocytes aging under normal conditions varied for different brain regions (Boisvert et al., 2018). The vulnerability of thalamic astrocytes in prion diseases could be attributed to intrinsic tropism of prions to this region, since the thalamus is impaired at the early stages and is the most severely affected at the advanced stage of prion diseases (Sandberg et al., 2014; Carroll et al., 2016; Makarava et al., 2020b). In 5XFAD mice, cortex and hippocampus are the first regions to deposit Aβ plaques and show signs of astrogliosis at younger ages (Oakley et al., 2006). However by 14 months of age, the thalamus exhibited the highest load of Aβ plaques in 5XFAD mice as judged by the uptake of an amyloid-specific tracer (Frost et al., 2020). In the current work, thalamic astrocytes reacted not only to the insults that are known to target thalamus, but also to injuries delivered to other brain regions. In fact, among four regions, thalamus showed the highest GSA scores with respect to both astrocyte reactivity and function in the TBI group (Fig. 6B). These results are consistent with previous clinical work that employed positron emission tomography ligand for measuring glia activation in humans. Up to 17 years after severe TBI, individuals showed profound chronic neuroinflammation in the thalamus, which was attributed to damages in thalamo-cortical tract (Ramlackhansingh et al., 2011; Scott et al., 2015). Moreover, neuroinflammation in thalamus has been proposed to serve as a marker of cortical injury and subsequent long-term cognitive deficits (Necula et al., 2021). In the Aged group, thalamus showed the highest GSA scores for the functional gene sets (Fig. 6B). Since the thalamus receives reciprocal projections from the entire cerebral cortex, its vulnerability could play an important role in understanding pathology and predicting disease outcomes (Grossman and Inglese, 2015; Scott et al., 2015).

Several intrinsic limitations of the current work are worth discussing. First, examining transcriptome of bulk tissues does not report on the actual heterogeneity in cell reactive phenotypes within individual brain regions or the diversity of cell responses to individual insults. Second, while the panel consists of genes that were selected to report on astrocyte function or reactive state, the contributions of other cell types cannot be excluded, especially under pathological conditions. Third, analysis of transcriptome on its own is not sufficient for defining functional phenotypes of the reactive state or making conclusions regarding the net neurotoxic or neuroprotective impact of astrocyte transformation. Moreover, region-specific differences in severity of astrocyte response can be interpreted in several alternative ways – for instance, as differences in intrinsic vulnerability of astrocytes to insults, or as differences in the strength of physiological response, whether maladaptive or not, to restore homeostasis. The current data should not be used for tracking the dynamics of individual genes, but rather for profiling of region-specific response. Sex-specific differences were previously found to influence glia reactivity in neurodegeneration associated with TBI and Alzheimer’s disease (Villapol et al., 2017; Guillot-Sestier et al., 2021), however, they were not subject of the current work. Considering the results and above limitations, the current work supports the idea that rigorous analyses of astrocyte transcriptome on a single cell level requires knowledge about the regional identity of astrocytes, especially for comparing the data acquired by independent studies.

Despite limitations, approaches that involve bulk tissue analyses might be well positioned to provide a holistic view of changes in astrocyte phenotype. Astrocytes translate mRNAs in their peripheral processes that are adjacent to synapses and enriched with mRNAs associated with biological functions including regulators of synaptic plasticity, GABA, glutamate metabolism and lipid synthesis (Sakers et al., 2017). One astrocyte can contact up to 100,000 synapses (Oberheim et al., 2009). Due to significant losses of peripheral astrocyte processes during isolation, the approaches that rely on isolation are likely to underestimate changes in gene sets that report on astrocyte functions. Recent work that employed singlenucleus RNAseq revealed a subpopulation of astrocytes in aged wild type mice and human brain that was phenotypically similar to a subpopulation of disease-associated astrocytes identified in the 5XFAD AD mouse model (Habib et al., 2020). While the approach in the current study lacked resolution of a singlenucleus RNAseq, we also observed considerable overlap in clusters corresponding to the same mouse model of AD and 24-month-old aged wild type mice. Remarkably, our study arrived at similar conclusions despite intrinsic limitations, such as a limited resolution due to analysis of only certain genes in bulk tissue. Nevertheless, the current approach could be useful for region-specific profiling of reactive phenotypes across multiple insults in humans.

## Author contributions

Conceptualization, I.V.B. and N.M.; Methodology, N.M., R.J.H. and N.T.; Investigation, N.M., O.M., J.C.C., R.J.H. and N.T.; Writing – Original Draft, I.V.B. and N.M.; Writing – Review & Editing, V.G., J.M.S. and D.L.; Funding Acquisition, I.V.B. and V.G.; Resources, K.M.; Supervision, I.V.B., V.G., J.M.S., and D.L.

## Supplementary material

### Supplemental Figures

**Figure S1. Co-immunostaining for Iba1 and GFAP showing activated microglia and pronounced astrogliosis in brains affected by neurodegeneration. (A)** Thalamus of prion-infected mice. **(B)** Normal thalamus. **(C)** Hippocampus of prion-infected mice. **(D)** Normal hippocampus. **(E)** Higher magnification image of prion-infected hippocampus. **(F)** Higher magnification image of normal hippocampus. **(G)** Cortex of 5XFAD mice. **(H)** Higher magnification image of 5XFAD cortex. Scale bar is 60 μm for A, B, C, D and G, and 20 μm for E, F and H.

**Figure S2. Ordered heat maps of gene expression in individual samples shown for four brain regions.**

**(A)** Ordered heat map for TBI and controls (Ctrl) visualizing the extent of global changes in gene expression at 7 and 60 days post insult in ipsilateral (ipsi) and contralateral (contra) hemispheres. Hypothalamus samples (HTh) were prepared from whole hypothalamus containing both ipsilateral and contralateral parts.

**(B)** Ordered heat map for MCAO and Sham visualizing the extent of global changes as well as variations between individual samples at 24 hours and 5 days post insult. **(C)** Ordered heat map for 5XFAD and wild type (WT) littermates. Females (F) and males (M) shown separately. **(D)** Ordered heat map for Aged mice (24-month-old) and control mice of various ages. Detailed description of sample groups is available in Table S1.

**Figure S3. Clustering of control samples occurs accordingly to brain region.** Clustering was performed using genes related to BBB regulation, lipid and energy metabolism, extracellular matrix, junction, myelination, channels, transporters, gliotransmitters and neurotransmitters, neuroprotection and neurotoxicity, and astrocyte-specific markers (Table S2).

**Figure S4. Brain region specificity of clustering is maintained after insults.** Grouped samples clustered accordingly to brain regions. The only condition with somewhat disturbed clustering is MCAO at 5 days post insult (MCAO-5d), which displayed substantial variability within individual samples. TBI is represented by ipsilateral Ctx, Hp and Th, and whole HTh. Ctx – cortex, Hp – hippocampus, Th – thalamus, HTh – hypothalamus.

**Figure S5. Relative changes in gene sets characteristic of astrocyte reactivity and function upon different insults.** Clustered heat maps of directed global significance scores for selected sample groups presented in four brain regions. TBI-7d is represented by ipsilateral Ctx, Hp and Th, and whole HTh. Ctx – cortex, Hp – hippocampus, Th – thalamus, HTh – hypothalamus.

**Figure S6. Examples of insult-specific changes in individual genes in cortex.** Plots of average normalized counts show individual sample values and standard deviation. Number of samples in each group is provided in Table S1. The statistical significance of changes in gene expression was calculated as part of a differential expression analysis. Significant *p*-values adjusted for false discovery rate are provided in Table S3.

### Supplemental Tables

**Table S1. Description of animal groups.**

**Table S2. Normalized counts for all individual animals, reported for four brain regions.**

**Table S3. Lists of differentially expressed genes (DEGs).** The table includes data for four brain regions (linear fold change ≥±1.2, adjusted *p<0.1).*

**Table S4. Lists of differentially expressed genes (DEGs) common between Aged and other groups.** The table includes data for four brain regions (linear fold change ≥±1.2, adjusted *p<0.1*).

