## Supplementary figures and images for "Region-specific homeostatic identity of astrocytes is essential for defining their reactive phenotypes following pathological insults"

### Figure S1

Figure S1

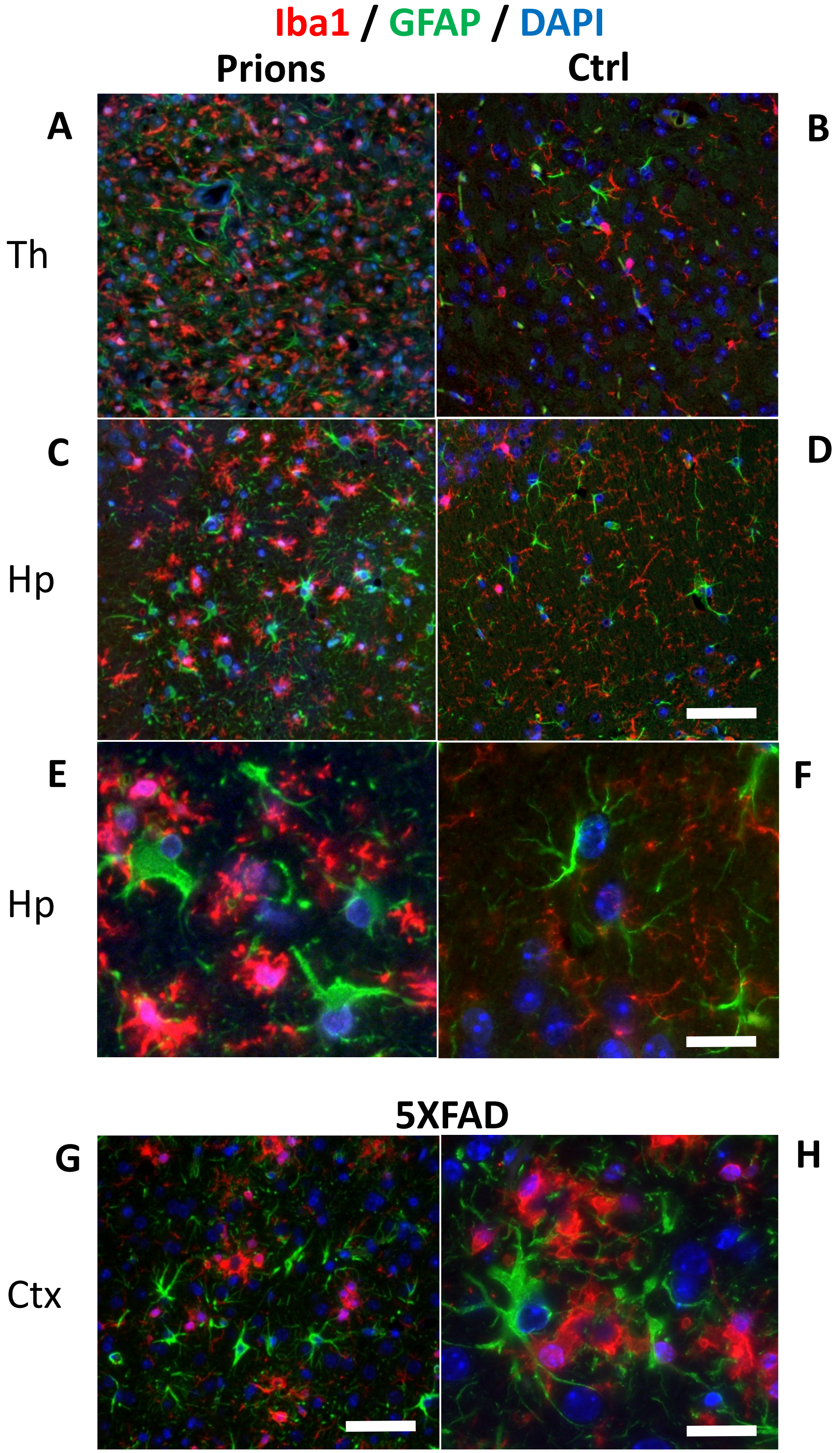

### Figure S2

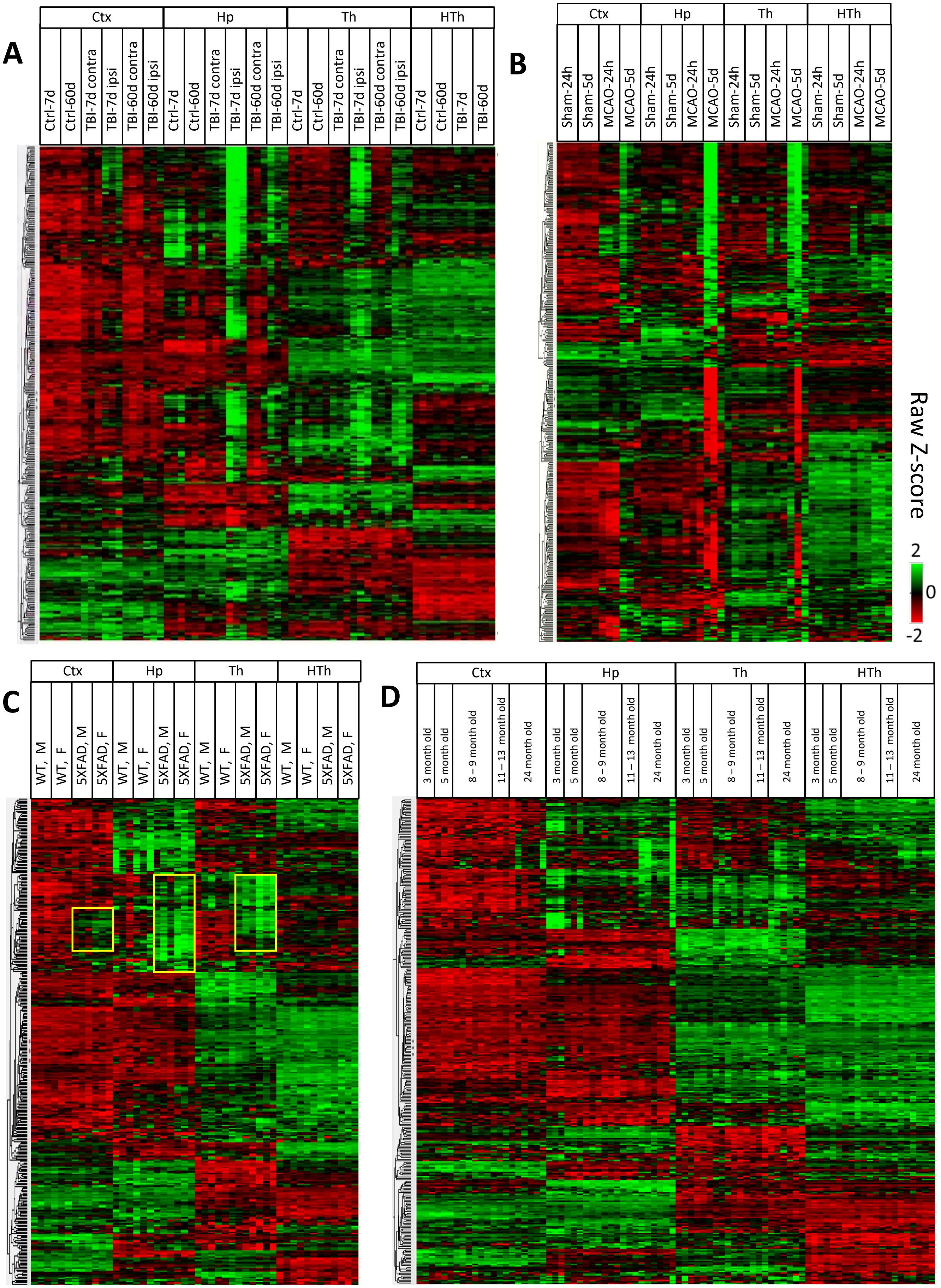

### Figure S3

Figure S3

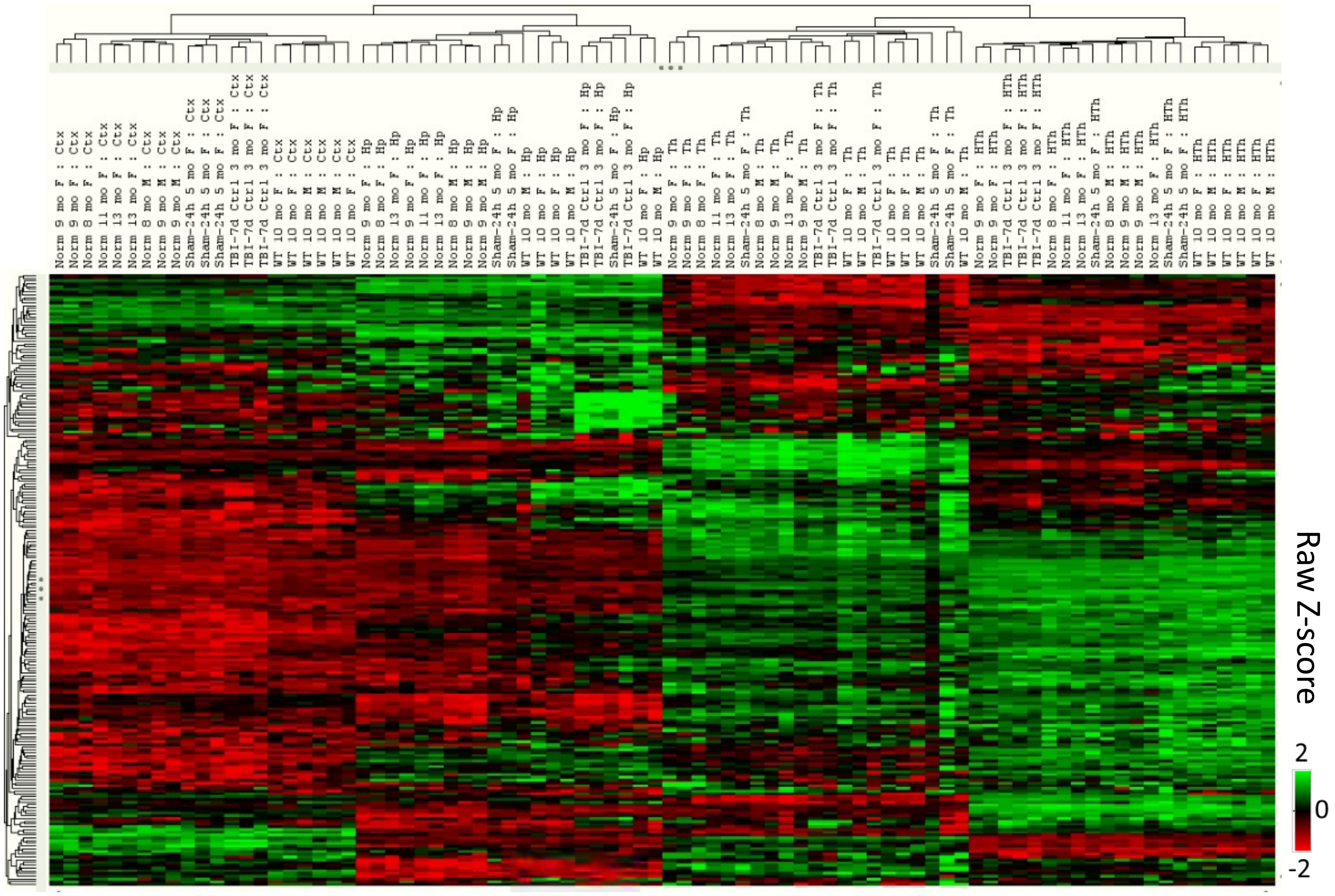

### Figure S4

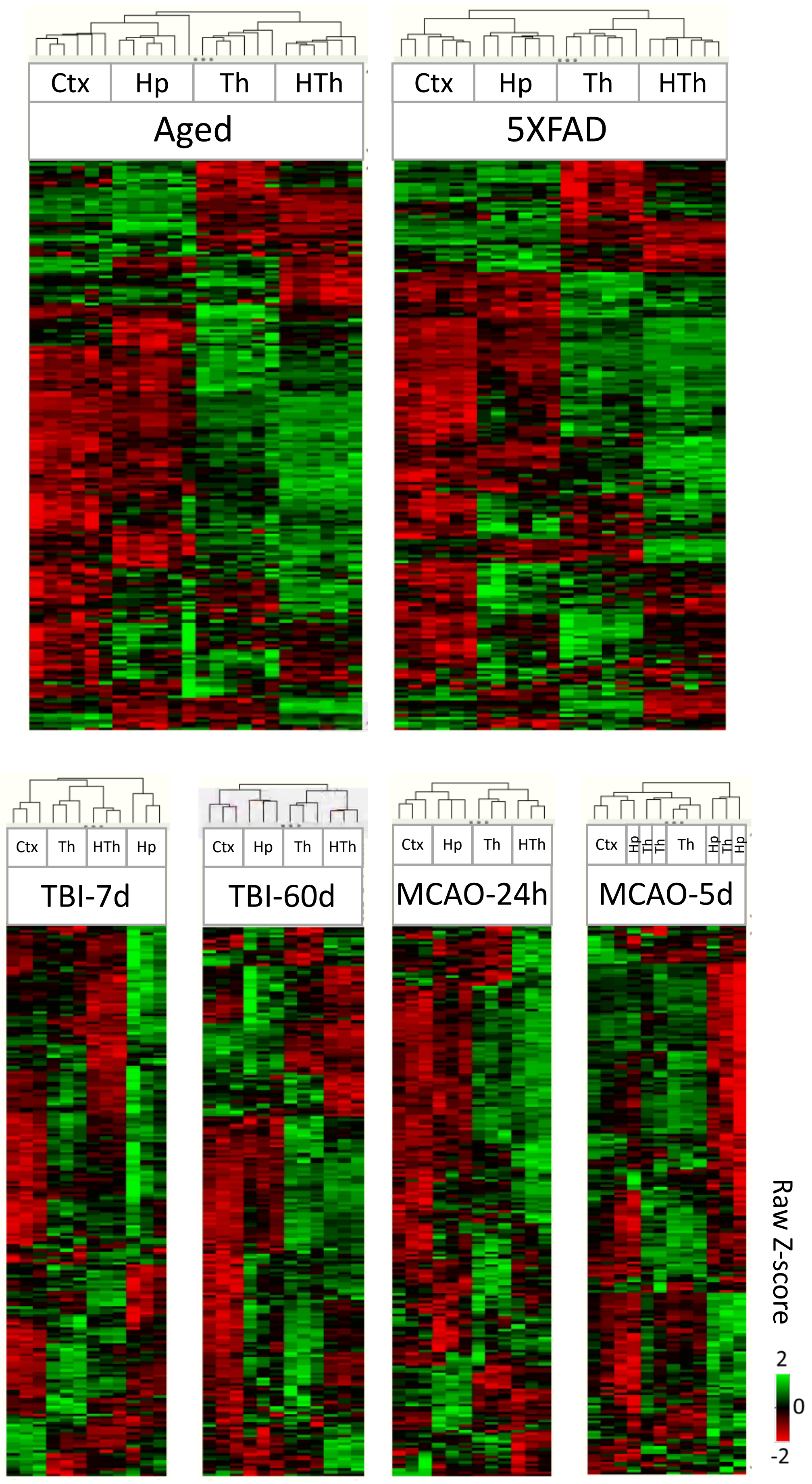

### Figure S5

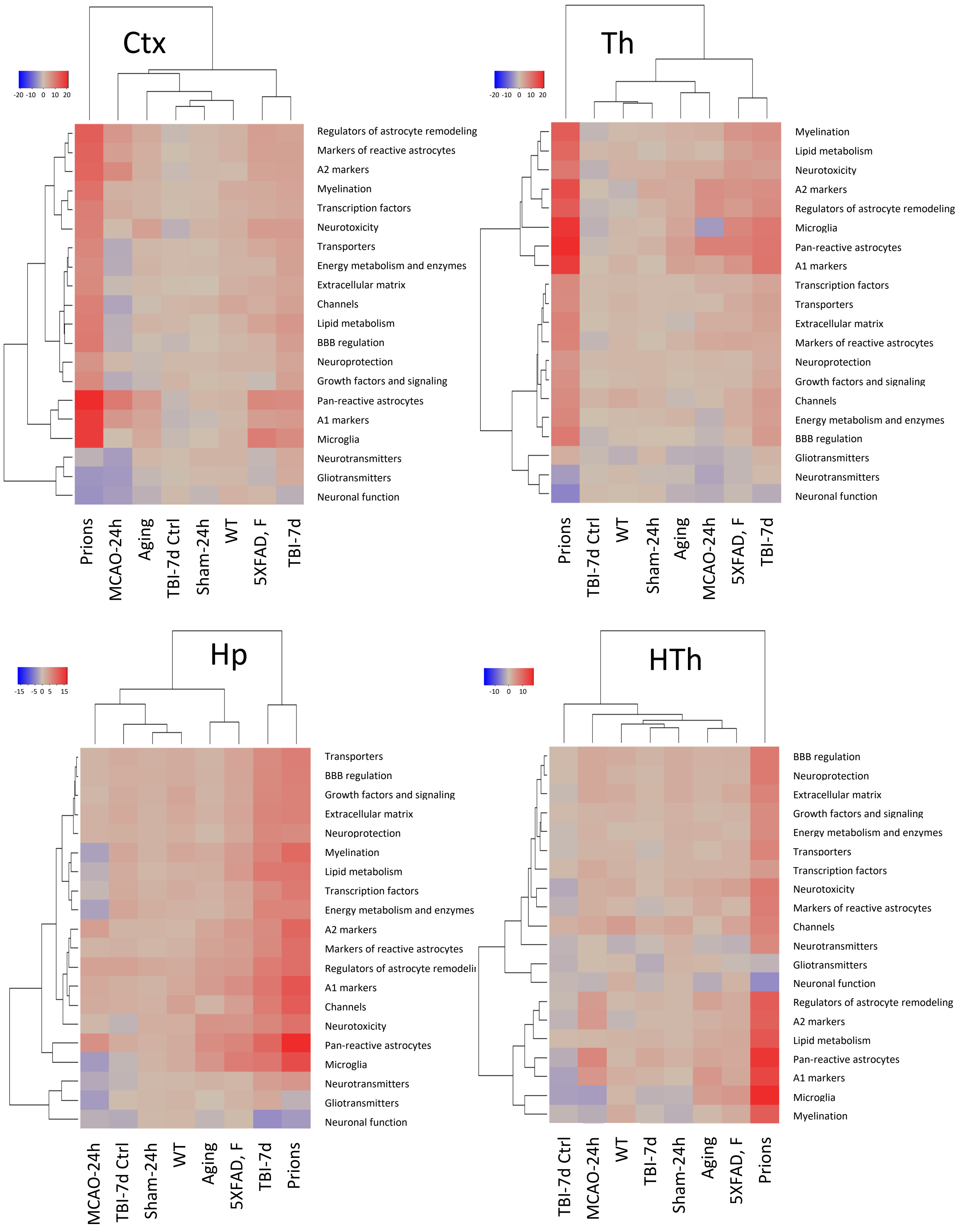

### Figure S6

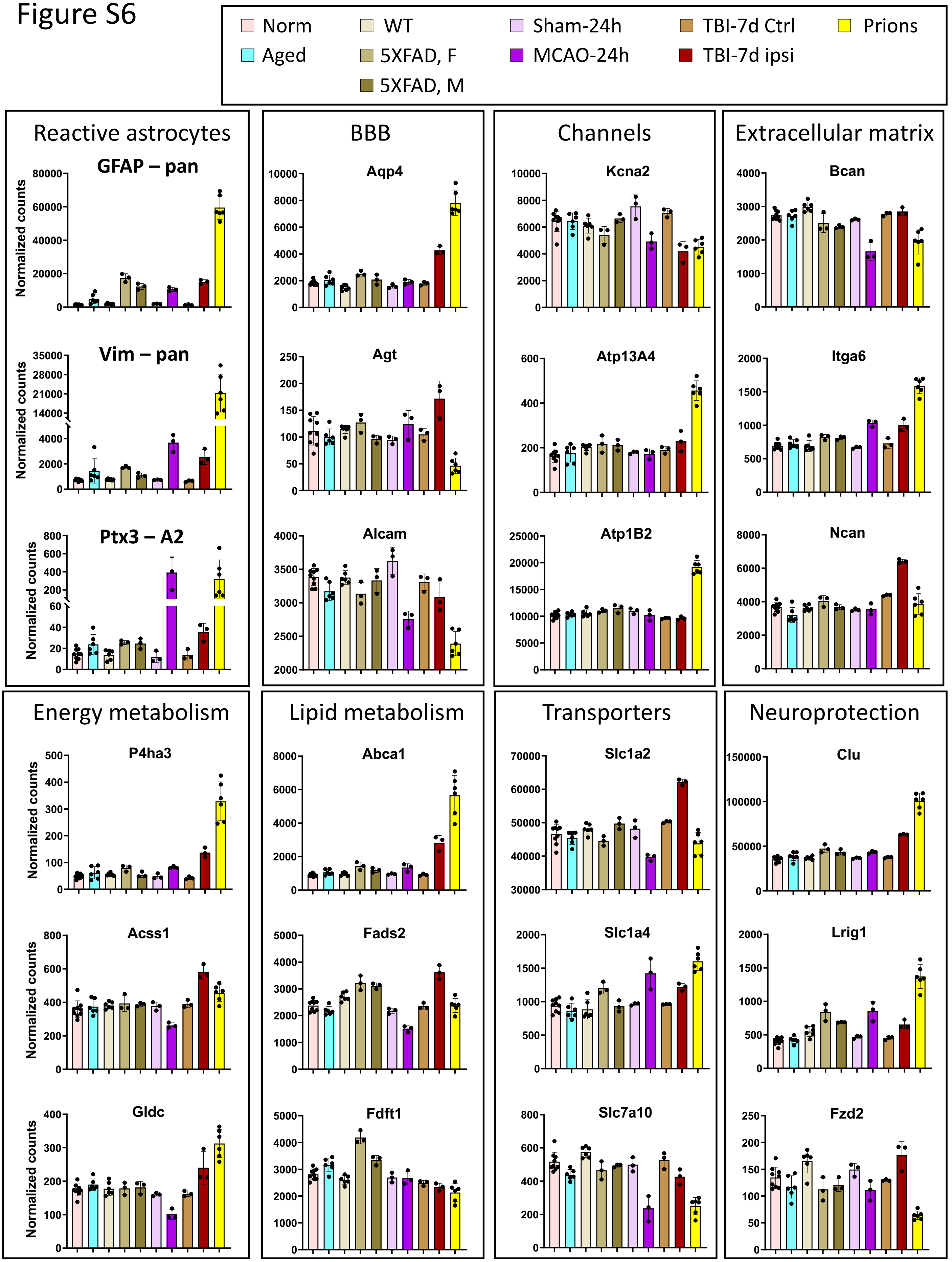
