## Supplementary material for "Region-specific homeostatic identity of astrocytes is essential for defining their reactive phenotypes following pathological insults": Table S1

**Table S1. List of animal groups analyzed using Astrocyte panel**

| Group name | Mouse strain | n | Age at insult | Insult | Time of euthanasia |
|---|---|---|---|---|---|
| Prions | C57Bl/6J | 4F+2M[1] | 6 weeks | 1% SSLOW i.p. | Terminal stage, dpi: F – 166, 166, 173, 173 [2]<br>M – 185, 175 (8 -13 month old) |
| Normal (Norm) | C57Bl/6J | 6F+3M | 6 weeks | 1xPBS i.p. | dpi: F – 197, 223, 223, 295, 346, 363; M – 203, 225, 229 (8 -13 month old) |
| TBI-7d ipsilateral | C57Bl/6J | 3F | 12 weeks | Left hemi CCI [3] | 7 dpi (3 month old) |
| TBI-7d contralateral | | | | | |
| TBI-60d ipsilateral | C57Bl/6J | 3F | 12 weeks | Left hemi CCI | 60 dpi (5 month old) |
| TBI-60d contralateral | | | | | |
| TBI-7d Ctrl | C57Bl/6J | 3F | n/a | Sham/no CCI | 3 month old |
| TBI-60d Ctrl | C57Bl/6J | 3F | n/a | Sham/no CCI | 5 month old |
| MCAO-24h | C57Bl/6J | 3F | 5 month | 60 min MCAO [4] | 1 dpi |
| MCAO-5d | C57Bl/6J | 3F | 5 month | 60 min MCAO | 5 dpi |
| Sham-24h | C57Bl/6J | 3F | 5 month | Sham | 1 dpi |
| Sham-5d | C57Bl/6J | 3F | 5 month | Sham | 5 dpi |
| Aged | C57Bl/6J | 3F+3M | n/a | Aging | 24 month old |
| 5XFAD | B6SJL | 3F+3M | n/a | 5 FAD [5] mutations | 10 month old |
| WT | B6SJL | 3F+3M | n/a | No mutations | 10 month old |

[1] F- females, M-males
[2] dpi – days past insult
[3] CCI – Controlled Cortical Impact
[4] MCAO – Middle Cerebral Artery Occlusion
[5] FAD – Familiar Alzheimer’s Disease
